# Segmented regression to streamline semi-mechanistic pharmacodynamic modelling of antibiotic effects

**DOI:** 10.64898/2026.11.03.763865

**Authors:** Anh Duc Pham, Wisse van Os, Jorn Brink, J. G. Coen van Hasselt, Linda B.S. Aulin

## Abstract

**Purpose:** Semi-mechanistic pharmacodynamic (PD) models quantify bacterial dynamics from time-kill (TK) assays, often showing multi-phasic kill and regrowth dynamics. Regrowth is typically captured with two subpopulations with distinct concentration-effect relationships. Since only total bacterial counts are observed, subpopulation dynamics must be inferred, causing identifiability issues, sensitivity to initial estimates, and model-testing burden. We aimed to establish an efficient and robust approach for developing semi-mechanistic PD models, demonstrated with 24-hour TK data for colistin against *Pseudomonas aeruginosa*.

**Methods:** A three-step workflow was developed: (1) log-transformed TK data were segmented into kill and regrowth phases, with linear regression estimating phase-specific slopes per concentration and replicate; (2) empirical slopes were pooled to derive concentration-effect relationships; (3) selected relationships and parameter estimates were incorporated into a semi-mechanistic PD model.

**Results:** Segmented regression captured kill and regrowth slopes at all concentrations. Slope saturation with increasing colistin and consistently weaker drug effect during regrowth indicated a less susceptible subpopulation, motivating a two-subpopulation model. The semi-mechanistic model adequately described bacterial dynamics at all concentrations, with parameters comparable to Step 2.

**Conclusion:** This three-step workflow enables efficient and robust identification of concentration-effect relationships, reducing model testing and computational burden for streamlined semi-mechanistic antibiotic PD model development.

## Introduction

PK/PD (pharmacokinetic/pharmacodynamic) principles play a fundamental role in optimizing antibiotic therapy by establishing dosing regimens that ma:imize efficacy while minimizing toxicity and the emergence of resistance(1). Accordingly, PK/PD evaluation is a key regulatory requirement in the development of new antimicrobial agents (2). PK/PD-guided dosing has been associated with reduced treatment costs, improved clinical outcomes and patient safety, and stabilization of antimicrobial resistance (1,3). Central to this framework are semi-mechanistic PD models, which integrate bacterial growth dynamics, antimicrobial killing, and resistance development, and can be applied across a wide range of bacterial infection models (4).

Semi-mechanistic PD models of antibiotics are commonly informed by *in vitro* static time-kill (STK) experiments (5), in which longitudinal bacterial dynamics in response to antibiotics are characterized. These experiments provide population-level observations that in combination with modelling generates quantitative and semi-mechanistic insight into antibacterial PD (4). The PD models incorporate and quantify key biological factors (e.g growth, drug effect, resistance states). This approach can separate bacteria specific parameters (growth, carrying capacity) from drug specific parameters (E_max_, EC_50_).

The development of semi-mechanistic PD models typically involves selection of several model components to capture distinct parts of observed bacterial growth dynamics and associated antibacterial concentration-effect relationships (6). Such components can for example capture initial lag in growth or bacterial regrowth associated with resistance or tolerance (6–8). Depending on the observed growth dynamics, a single or, more often, more often multiple bacterial subpopulations can be included (7,8). Such subpopulations are inferred from the different phases often observed in the STK curves, e.g. rapid initial kill, then slower kill or regrowth can often be represented by a two-population bacterial model. Generally, the choice of model components should be anchored in the mechanistic understanding of the system while being guided by the data, including pre-excising or *de novo* development of resistance. Similarly, multiple concentration-effect relationships can be considered (e.g., linear, E_max_, sigmoidal), and these relationships can differ between bacterial subpopulations (9,10). Although an effect function with a plateau is often the most biologically plausible, the choice of which relationship to implement is data driven. Only the observed effects within the studied sampling time and concentration range can inform parameter estimation, leading to practical identifiability issues when trying to estimate an unobserved ma:imum effect (11–13).

In practice, model structures of semi-mechanistic PD models for antibiotics often involve an iterative process of considering many possible model structures and components. To illustrate, one of the most widely used resistance model structure is a two-population model with a first-order transition from a susceptible (*S*) to a resistant (*R*) population (6). As STK assays often only quantify the total population as colony forming units (CFU)/mL, both subpopulations and their distinct behaviour must be inferred. The limited information in TK data combined with the complexity of semi-mechanistic models can often lead to practical identifiability issues, parameter bias, and high sensitivity to initial estimates (11–13). Since subpopulations may also differ in their concentration-effect relationships, several combinations often need to be evaluated, making the final model depend on the sequence of modelling decisions.

This work aims to address some of the aforementioned challenges by proposing a workflow to support the efficient development of antibiotic semi-mechanistic PD models for STK data. The workflow constitutes a three-step process wherein phase-specific slopes are estimated from observed STK data, subsequently used to fit concentration-effect relationships, and finally integrated into a semi-mechanistic PD model. We demonstrate this workflow using STK data of colistin against a *Pseudomonas aeruginosa* strain.

## Methods

### Bacterial strains and antibiotics

A clinical *P. aeruginosa* isolate, PA11976, was used throughout the study. Bacteria were directly cultured from cryostocks on agar plates. The plates were incubated overnight and then harvested and diluted to the desired inoculation concentration (10^6^ CFU/mL). An aqueous colistin stock solution of 10 mg/mL was prepared by dissolving colistin sulphate (Cayman, USA) in deionized water followed by filter sterilization.

MIC was determined using a microbroth dilution method. Bacterial were recovered from cryostocks, streaked on Mueller-Hinton (MH) agar plates and incubated overnight. Bacterial cells were first harvested to prepare 0.5 McFarland standards in PBS (∼10^8^ CFU/mL), followed by dilution to achieve a target inoculum of ∼10^5^ CFU/mL in 96-well plates. Experiment was carried at 37°C for 18 to 20 hour period.

### Time-kill experiments

The TK experiments were conducted in duplicate in culture tubes with 25 mL cation-adjusted MH broth (VWR, Belgium). Tubes were inoculated with an overnight culture to a density of 10^6^ CFU/mL and incubated at 37°C with continuous shaking at 200 rpm. Samples (1 mL) were collected at 0, 2, 4, 6,12 and 24 h post inoculation for bacterial quantification. For this, 100 µl of serially diluted liquid culture was plated on MH agar plates with bead plating and semi-automated CFU counting (aCOLyte, Synbiosis) (14).

### Workffow

A workflow was developed to enable fast robust characterization of TK data (**Figure 1**), consisting of three steps: (1) Segmented regression of kill and regrowth phases; (2) Concentration-effect relationship identification; (3) Semi-mechanistic PD model development.

**Figure 1.**
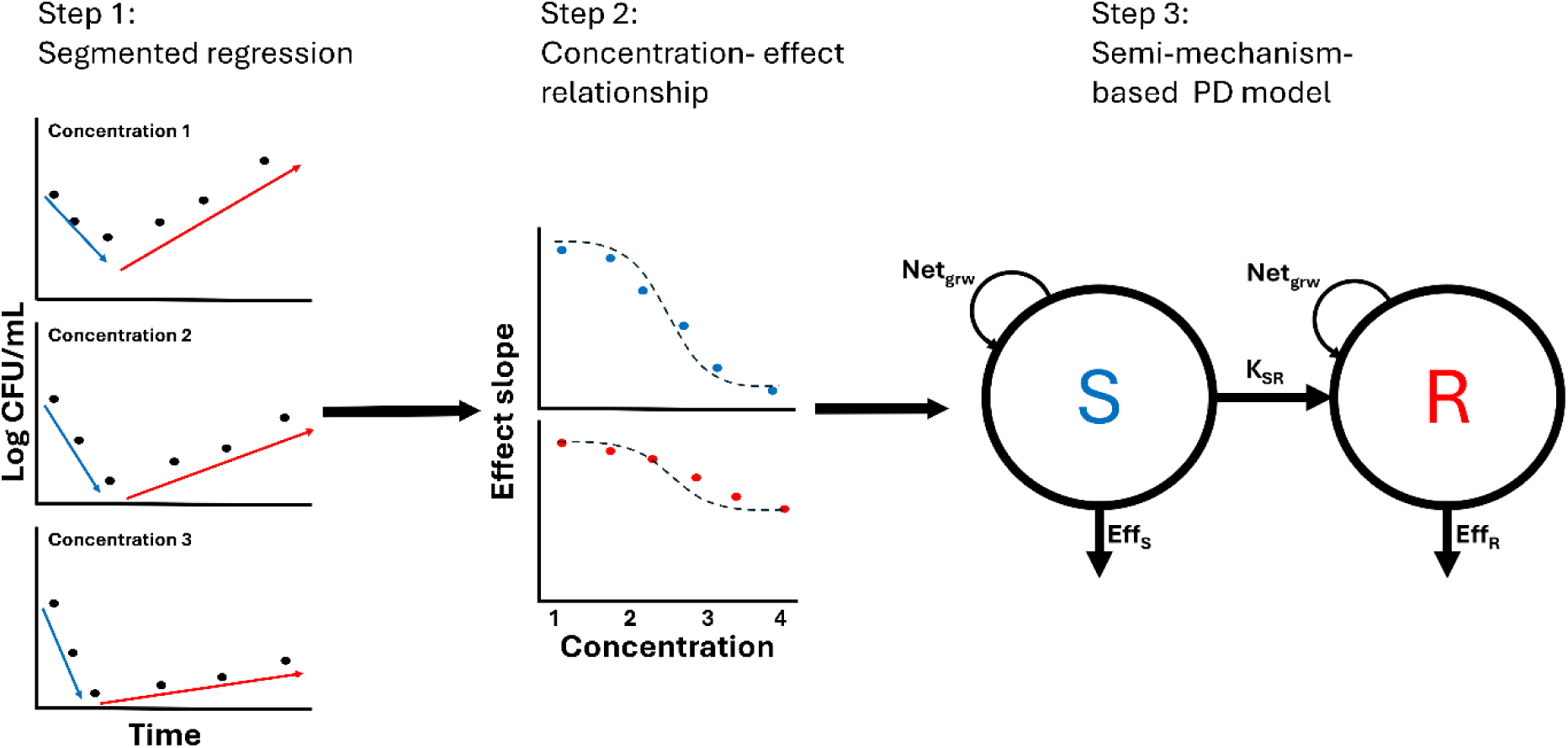
Proposed modelling workffow. Step 1: Segmented regression of initial kill phase (blue) and subsequent regrowth phase (red) to derive concentration-specific effect slopes. Step 2: Fitting separate concentration-effect relationships for kill and regrowth phase (dashed line). Step 3: Incorporating identified relationships into two-subpopulation semi-mechanistic-based PD model including a susceptible (S) and a resistant (R) subpopulation.

### Step 1: Segmented regression of kill and regrowth phase

An initial segmented regression-based approach was developed to enable fast, robust characterization of STK data. The data-driven method focuses on extracting features directly from observed bacterial dynamics rather than fitting semi-mechanistic models at an early stage.

The CFU values are first transformed to the natural logarithmic scale. For each antibiotic concentration, the STK profile is segmented into two phases. The CFU-declining phase represents drug-driven bacterial killing, typically observed from treatment initiation to the time of minimum CFU. Following initial kill, the CFU-increasing phase represents bacterial regrowth. The segmentation time point is determined empirically based on the time where the minimum CFU is observed (*t_min_*), which in the simplest case is consistent across concentrations. Alternatively, *t_min_* can be inferred using automated segmentation algorithms (15–17). Within each phase, bacterial dynamics are approximated using a linear regression model on the natural log scale, and separate slopes are estimated for the kill and regrowth phase at each concentration:

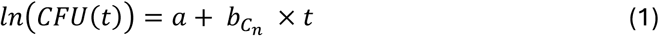

Here, *a* is the intercept, corresponding to *ln* (*CFU*(*t*_0_)) at t_0_, i.e. the inoculum or the estimated starting concentration, for the initial phase and *ln* (*CFU*(*t_min_*)) for the regrowth phase at t_min_, and *b_Cn_* is the concentration-specific slope (in ln (*CFU*/*mL*)/ℎ) representing the net rate of bacterial kill or growth for a specific concentration *C_n_*. CFU observations and model predictions are log-transformed to stabilize variance. The lower limit of detection (LLOD) was set to 10 CFU/mL. Observations below LLOD are imputed as half of LLOD.

Two patterns that can commonly be present in the STK data are a growth lag and a stationary phase. These phases should be excluded from Step 1, as these data do not inform concentration-effect relationships.

### Step 2: Concentration-effect relationship identification

The second step allows for the identification of the most suitable concentration-effect relationships for the kill and regrowth phase, respectively. The estimated concentration-specific effect slopes (*b_c_*_0,_*b_c_*_1_, *b_c_*_2_… *b_cn_*) from Step 1 are combined per phase to derive relationships describing the growth or kill rate over the antibiotic gradient. Three different concentration-effect relationships are evaluated per phase to describe the net effect (*Net_eff_*) including growth (*b_c_*_0_) and antibacterial effect (*f_eff_*); linear (Eq. 2), E_max_ (Eq. 3 v=1) or sigmoidal E_max_ (Eq.3 v≠1)

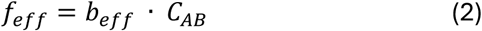

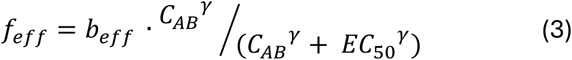

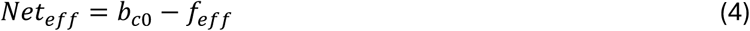

where *b_c_*_0_ represents the bacterial growth rate (ln(CFU) per unit of time) in the absence of drug, *C_AB_* is the antibiotic concentration, *b_eff_* represents either the slope (Eq. 2) or the ma:imum effect (Eq.3), *EC*_50_ is the concentration producing half-ma:imal effect, and *γγ* is the Hill coefficient.

Each concentration-effect relationship is fitted to the regression-derived slopes over concentration, and model performance is compared by visual inspection of model fit, parameter precision and using the Akaike Information Criterion (AIC), where the lowest AIC is preferred. This step enables rapid identification of the most suitable concentration-effect relationships and informs the initial estimates when re-estimated in the full mechanism-based PD model constructed with ordinary differential equations (ODEs).

### Step 3: Semi-mechanistic PD model development

In Step 3, the derived concentration-effect relationships identified in Step 2 are implemented in a two-subpopulation semi-mechanistic PD model (Figure 1c) in a nonlinear mi:ed effects framework. In the model, the total bacterial burden (CFU) is defined as the sum of two bacterial subpopulations; susceptible bacteria (S) and resistant bacteria (R). These two subpopulations allow the model to capture biphasic killing and regrowth commonly observed in STK data. The bacterial subpopulations are assumed to share the same intrinsic growth behavior, governed by a logistic growth function:

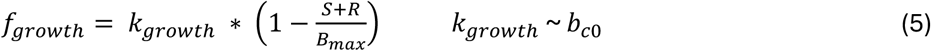

*k_ggroggt_*_ℎ_ is the bacterial growth rate constant and *B_max_* represents the system’s carrying capacity. The initial estimate of *k_ggroggt_*_ℎ_ was based on Step 2 as *b_c_*_0_. *B_max_* can be based on the observed control data from the TK experiment at 24h.

Drug-mediated killing is modelled according to the concentration-effect relationships identified in Step 2. The regression-derived parameters are transformed to the appropriate scale and are subsequently used to inform initial estimates. Specifically, the drug effect parameter *k_eff_*, representing either a slope or an E_max_ parameter, is approximated as *b_eff_*, where logarithmic transformation ensures consistency between regression-based and ODE models formulated on the normal scale. In the ODE models, drug-mediated killing is modelled as a concentration-dependent effect acting on each bacterial state where the killing phase informed drug effects on *S* and regrowth phase drug effects on *R*:

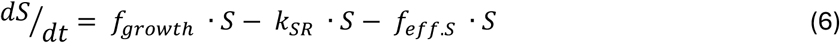

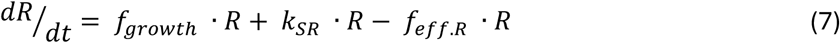

Where the drug effect function *f_eff_* is defined using concentration-effect relationships identified during Step 2 (Eq. 2 or 3) after transforming *b_eff_* to *k_eff_* as specified above and where *b_c_*_0_ is replaced by *f_ggroggt_*_ℎ_. For Eq. 3, EC_50_ and *γγ* are implemented without transformation. Time-dependent changes in population drug susceptibility are described by addition of an unidirectional transition from subpopulation *S* to *R*, mediated by the transition rate constant *k_SR_*. The rate constant describes the transition between the empirically segmented phases from previous steps, where the phases come to represent the bacterial subpopulations. At the start of the experiment, 100% of the bacterial population is assumed to belong to subpopulation *S*, and *k_SR_* is set to 0 for the drug-free condition. Residual unexplained variability was evaluated using additive error model over log10 transformation of the STK data.

### Software

Data visualisation, statistical analyses, and model evaluation were carried out in R 4.6 (18). Step 2 was performed using nonlinear least squares to estimate model parameters from the *stats* package v4.6 (18). The nonlinear mixed-effects modelling in Step 3 was performed using nlmixR 2 v5.0 (19) with first-order estimation.

## Results

We demonstrated the three-step workflow to a case study involving a STK experiment of colistin and one *P. aeruginosa* strain. The code for the proposed workflow and our proof-of-concept case study can be found here: https://github.com/LeidenPharmacology/SegmentedTK

### Step 1: Segmented regression analysis

The TK data are shown in **Figure 2** and **S1**). All profiles of drug exposed bacteria exhibited an initial kill phase. At low concentrations (≤0.5 µg/mL), killing was insufficient to suppress regrowth, whereas at higher concentrations no regrowth was observed. The inflection point between the killing phase and the regrowth phase (t_min_, identified as minimum observed CFU) was observed between 2 and 12h, with a mean of 4.2h (**Figure S1**). Kill and regrowth slopes were successfully derived for all concentrations (**Figure 2A**).

**Figure 2.**
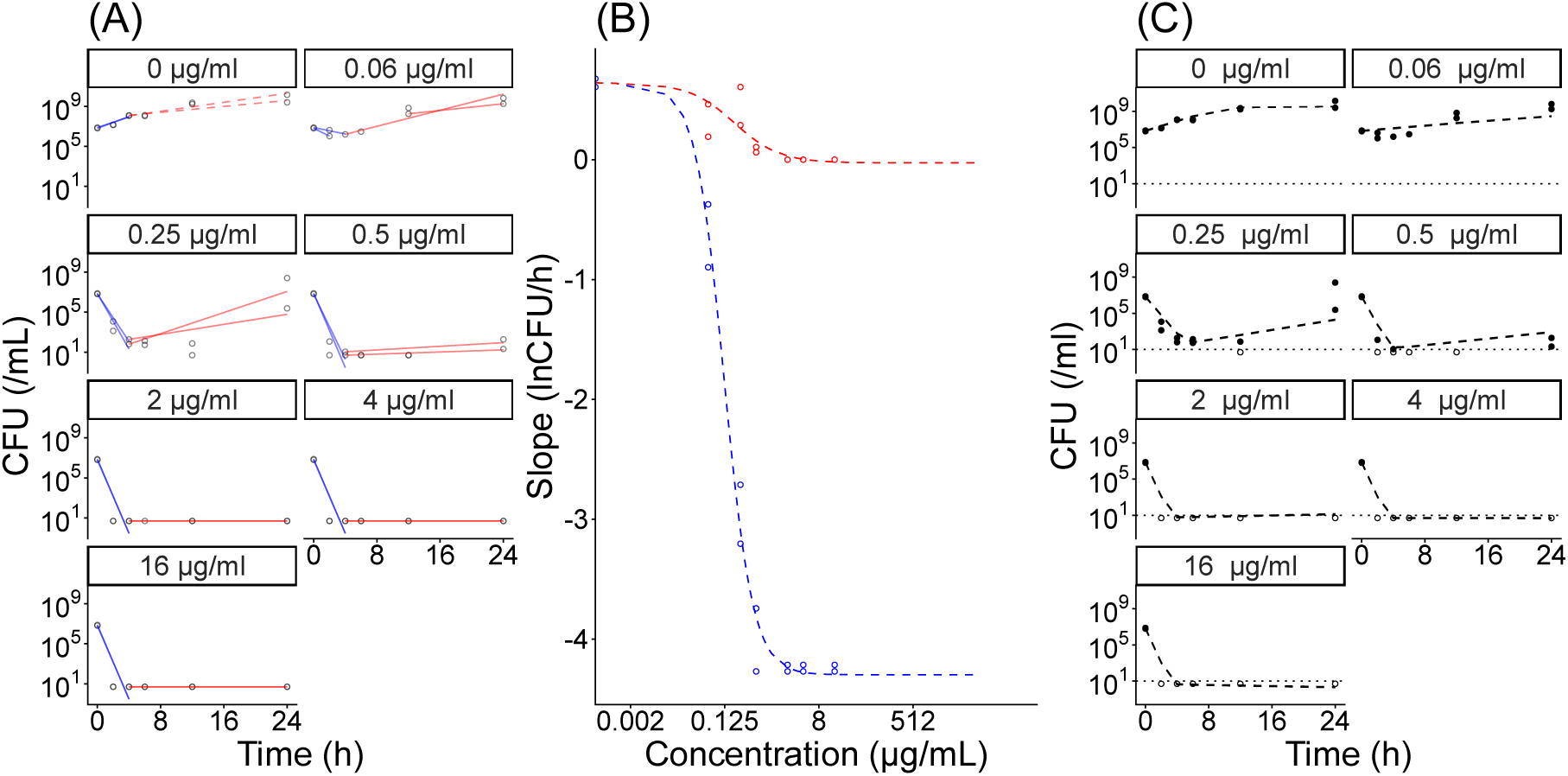
Three-step workflow for developing mechanism-based pharmacodynamic model of the effect of colistin on isolate PA11976 (**A**) Step 1: identified effect slope from time kill data (**B**) Step 2: Regression-derived concentration– effect relationships based on the slopes for each of the two segments. (**C**) Step 3: Semi-mechanistic pharmacodynamic model describing antibiotic effects and bacterial growth dynamics

### Step 2: Concentration-effect relationship characterization

The determined slopes were evaluated over the concentration gradient to determine phase-specific concentration-effect relationships. The initial slope of the drug-free condition was positive and provide the intrinsic underlying growth rate (0.64 LogCFU/ml/h), which was assumed to be the same for both phases. Kill-phase slopes became increasingly negative with rising antibiotic concentrations, demonstrating concentration-dependent killing, while higher drug levels also reduced slope values for the regrowth phase. The effect of increasing drug concentrations on the slopes plateaued at 2 µg/mL for both the killing and the regrowth phase, suggesting nonlinear concentration–effect relationships (**Figure 2B**). An overview of tested models is shown in **Table 1**, where the killing phase and regrowth phase were best represented by a sigmoidal E_max_ and a E_max_ model, respectively, as indicated by lowest AIC values (**Table 1**). Across concentration–effect relationships, the regrowth phase showed a reduced antibiotic susceptibility compared with the initial phase. The concentration–effect relationship exhibited reduced drug susceptibility during regrowth, indicated by lower slope (slope_kill_ 0.40 vs. slope_regrowth_ 0.05 h^-1^), E_max_ (E_max,kill_ 5.1 vs. E_max,regrowth_ 0.67 h^-1^), and sigmoidal E_max_ (E_max,kill_ 4.9 vs. E_max,regrowth_ 0.72 h^-1^) estimates. These results align with the biological interpretation of the regrowth phase representing less sensitive or resistant bacteria.

**Table 1:**
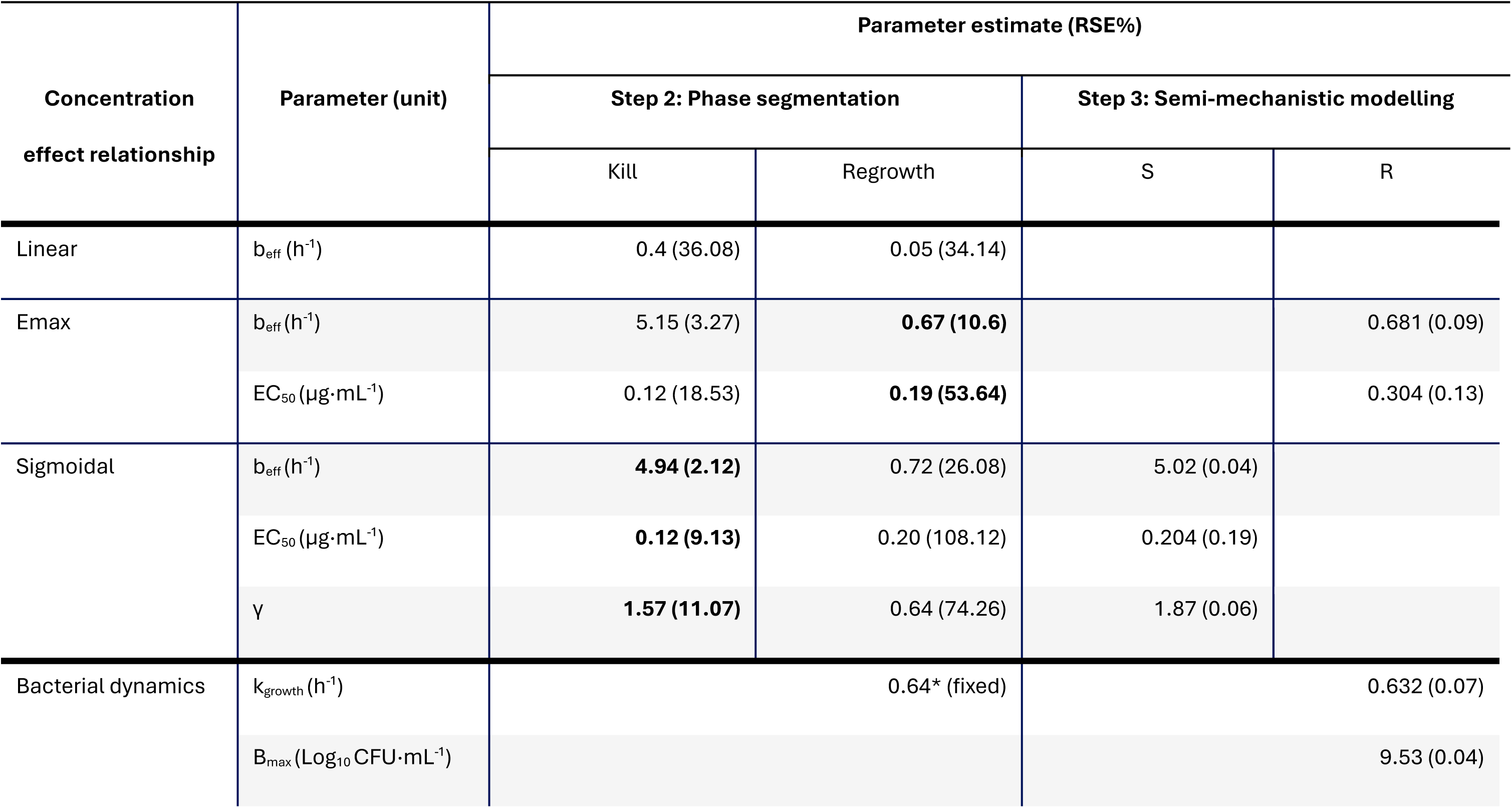
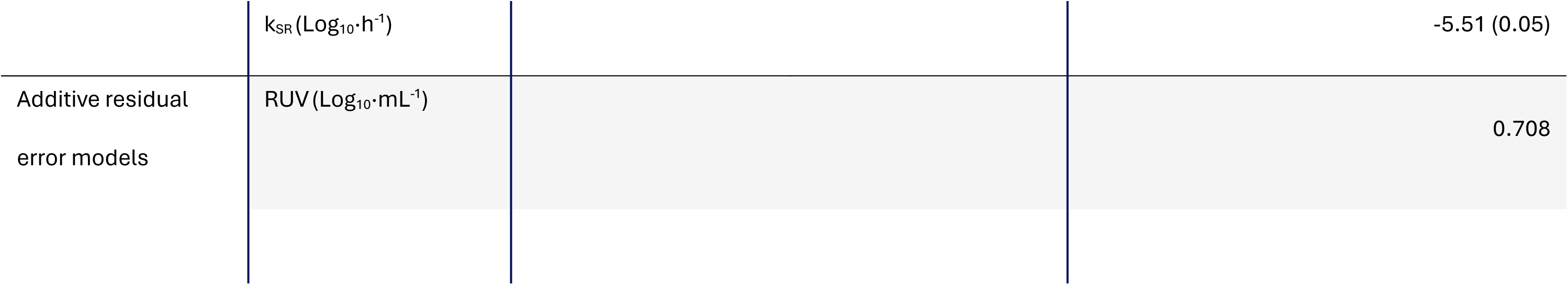
Model and parameter (RSE%) estimated by phase segmentation (Step 2) and mechanistic modelling (Step 3) for isolate PA11976. Selected concentration-effect relationships from Step 2 are indicated in bold and integrated in the mechanistic model in Step 3.

| **Concentration effect relationship** | **Parameter (unit)** | **Parameter estimate (RSE%)** |  |  |  |
|---|---|---|---|---|---|
|  |  | **Step 2: Phase segmentation** |  | **Step 3: Semi-mechanistic modelling** |  |
|  |  | Kill | Regrowth | S | R |
| Linear | b<sub>eff</sub> (h<sup>-1</sup>) | 0.4 (36.08) | 0.05 (34.14) |  |  |
| Emax | b<sub>eff</sub>(h<sup>-1</sup>) | 5.15 (3.27) | **0.67 (10.6)** |  | 0.681 (0.09) |
|  | EC<sub>50</sub> (µg·mL<sup>-1</sup>) | 0.12 (18.53) | **0.19 (53.64)** |  | 0.304 (0.13) |
| Sigmoidal | b<sub>eff</sub>(h<sup>-1</sup>) | **4.94 (2.12)** | 0.72 (26.08) | 5.02 (0.04) |  |
|  | EC<sub>50</sub> (µg·mL<sup>-1</sup>) | **0.12 (9.13)** | 0.20 (108.12) | 0.204 (0.19) |  |
|  | γ | **1.57 (11.07)** | 0.64 (74.26) | 1.87 (0.06) |  |
| Bacterial dynamics | k<sub>growth</sub> (h<sup>-1</sup>) | 0.64* (fixed) |  | 0.632 (0.07) |  |
|  | B<sub>max</sub> (Log<sub>10</sub> CFU·mL<sup>-1</sup>) |  |  | 9.53 (0.04) |  |
|  | k<sub>SR</sub> (Log<sub>10</sub>·h<sup>-1</sup>) |  |  | -5.51 (0.05) |  |
| Additive residual error models | RUV (Log<sub>10</sub>·mL<sup>-1</sup>) |  |  | 0.708 |  |

*Fixed to *b_c_*_0_from Step 1

AIC of Linear, E_max_ and sigmoid E_max_ are 66.36, 14.61, and 4.51 for S and 16.15, -8.06, and -6.84 for R, respectively. Parameters included are b_eff_ (slope or maximum kill rate), EC_50_ (concentration at half max kill rate), v (hill coefficient), k_growth_ (maximum growth rate), B_max_ (maximum population), k_SR_ (S to R phase transition rate), RUV (residual unexplained variability of growth/effect data represent additive residual SD)

### Step 3: Semi-mechanistic PD model

The concentration-effect relationships identified in Step 2 were implemented in the mechanism-based PD model where the kill phase informed the *S* subpopulation and the regrowth phase the *R* subpopulation. The final model could adequately describe the time course of bacterial growth, drug effects, and regrowth across all tested colistin concentrations (Fig 2c). The estimated parameters are comparable with those derived in Step 2 (Table 1). Initial rapid killing was captured by the susceptible subpopulation *S*, followed by delayed regrowth driven by the emergence of less-susceptible bacteria *R*. Although Step 2 highly informed the mechanism-based PD model, some model parameters are not informed by previous steps and need to be estimated in Step 3 without any guidance of initial estimates, e.g. *k_SR_*.

## Discussion

In this study, we propose a curve-segmentation-based workflow for aiding the development of semi-mechanistic PD models from STK data. This addresses the need for more efficient and robust approaches when developing complex semi-mechanistic bacterial PD models, where practical parameter identifiability, sensitivity to initial estimates, and dependence on the sequence of modelling decisions can make reproducible model development difficult (7,20,21). The workflow begins with estimating separate slopes in the STK data (i.e., for the killing and regrowth phases), which are then used to derive phase-specific concentration-effect relationships and inform the structure and initial parameter estimates of a semi-mechanistic PD model, providing insight into concentration-dependent dynamics. By characterising the distinct phases separately, the workflow reduces comple:ity while e:tracting informative features that can guide model building and pre-screening of concentration-effect relationships. These relationships are integrated into semi-mechanistic PD models, which are highly valuable tools to understand and improve antibiotic treatment and suppress resistance development. There is range of such semi-mechanistic models, each associated with different assumptions regarding the underlying mechanism of the observed regrowth. We illustrated the workflow using colistin and a *P. aeruginosa* isolate using a common two subpopulation with *de novo* resistance as proof-of-concept.

The bacterial kill and regrowth slopes estimated with the regression approach capture the key STK dynamics in a simple and robust way. This approach is similar to curve-stripping approaches commonly used in noncompartmental PK analysis (22) but, to our knowledge, has not been applied to bacterial PD data. It is important to note that this approach assumes exponential (linear on the ln CFU scale) killing and regrowth within each segmented phase, which is generally appropriate for bacterial dynamics where no lag or stationary phases are present. As our proof-of-concept dataset did not include a lag phase and stationary phase, such dynamics are not currently implemented in the workflow but could be considered in the future by expanding the phase segmentation to explicitly include such phases. In our case example, the regression-based estimates also provided reasonable initial values for the semi-mechanistic model, facilitating convergence and potentially improving parameter estimation.

The bacterial curves are segmented into phases where one of the subpopulations is assumed to be dominant, *S* in the kill phase and *R* in the regrowth phase, and the associated regression slopes are assumed to represent the behaviour of the dominant subpopulation. Such separation is commonly used to capture the regrowth of bacteria under antibiotic pressure (23). The breakpoint between the phases, or the point of segmentation, is empirically derived from the minimum observed CFU in our case example. Hence, transferring to semi-mechanistic model required the addition of a transition parameter. An alternative strategy to the empirically derived breakpoint would be to estimate the breakpoint as described in the Methods. Such an estimated parameter could inform the transition from *S* to *R*, often described using a transition rate in semi-mechanistic PD models, e.g. k_SR_. Our current workflow does not include this translation into model parameters. The choice of segmentation directly influences the derived phase-specific parameters and should therefore be considered carefully when applying the workflow. In addition, the stationary phase of control experiment can be used for driving estimation of the system carrying capacity B_max_ while the observed inoculum can either be estimated or fi:ed from the data.

We have demonstrated the utility of the workflow with a single-strain case example using *P. aeruginosa* and colistin. As a next step it should be applied and validated across more antibiotics and bacterial species. Additionally, we applied the workflow of one of the most common semi-mechanistic PD model structures describing bacterial dynamics, including regrowth. There are alternative model structures with different underlying assumptions for which our workflow could also be used. This could include models incorporating adaptive or preexisting resistance (23–25), where the derived slopes would analogously inform the associated concentration-effect relationships, although the approach would have to be slightly modified. In particular, adaptive resistance may require a shared concentration-effect relationship, where the kill phase informs the starting sensitivity and the regrowth phase the sensitivity after adaptive resistance rather than two separated subpopulations. Pre-existing resistance subpopulation or population heterogeneity of pathogen are of relevance to our case example as such behaviour has been reported for colistin previously (26), which might make a preexisting subpopulation without transitions an alternative model that could be applied in this case. However, the aim of this paper is not to identify the optimal semi-mechanistic model structure but support the identification and incorporation of concentration-effect relationships. The workflow could also be applied to a multi-strain context, where pathogen-level behaviour can be characterised and variability between strains can be quantified. More broadly, the pre-screening step may help identify informative concentration ranges, supporting adaptive and efficient e:perimental design.

In this work, we have proposed a workflow that makes the development of mechanism-based PD models for antibiotics faster and more robust. Faster, more reliable PD models are important to support dosing optimisation and antimicrobial resistance mitigation, highlighting the practical value of this workflow beyond methodological convenience.

## Declarations

### Funding and Confficts of interests

This project has received funding from the European Union’s Horizon 2020 research and innovation programme under the Marie Skłodowska-Curie grant agreement No 861323. The authors have no conflict of interest to declare.

## Abbreviations

PK: pharmacokinetic
PD: pharmacodynamic
STK: static time-kill
CFU: colony forming units
*S*: susceptible
*R*: resistant
MH: Mueller-Hinton
LLOD: lower limit of detection

## References

1. Rodríguez-Gascón A, Solinís MÁ, Isla A. The Role of PK/PD Analysis in the Development and Evaluation of Antimicrobials. Pharmaceutics. 2021 Jun 1;13(6):833. doi:10.3390/PHARMACEUTICS13060833 PubMed PMID: 34205113.

2. European Medicines Agency. Guideline on the use of pharmacokinetics and pharmacodynamics in the development of antimicrobial medicinal products [Internet]. 2016 [cited 2026 Jul 26]. Available from: https://www.ema.europa.eu/en/use-pharmacokinetics-pharmacodynamics-development-antibacterial-medicinal-products-scientific-guideline

3. Dubey V, Reza N, Hope W. Antimicrobial pharmacodynamics: defining regimens of antibiotics that are safe, effective and prevent the emergence of resistance. Philos Trans R Soc Lond B Biol Sci. 2026 Apr 30;381(1949). doi:10.1098/RSTB.2024.0340/481513 PubMed PMID: 42057718.

4. Nielsen EI, Friberg LE. Pharmacokinetic-pharmacodynamic modeling of antibacterial drugs. Pharmacol Rev. 2013 Jul;65(3):1053–90. doi:10.1124/PR.111.005769 PubMed PMID: 23803529.

5. Nielsen EI, Viberg A, Löwdin E, Cars O, Karlsson MO, Sandström M. Semimechanistic Pharmacokinetic/Pharmacodynamic Model for Assessment of Activity of Antibacterial Agents from Time-Kill Curve Experiments. Antimicrob Agents Chemother. 2007 Jan;51(1):128–36. doi:10.1128/AAC.00604-06 PubMed PMID: 17060524.

6. Minichmayr IK, Aranzana-Climent V, Friberg LE. Pharmacokinetic/pharmacodynamic models for time courses of antibiotic effects. Int J Antimicrob Agents. 2022 Sep 1;60(3). doi:10.1016/J.IJANTIMICAG.2022.106616 PubMed PMID: 35691605.

7. Jacobs M, Grégoire N, Couet W, Bulitta JB. Distinguishing Antimicrobial Models with Different Resistance Mechanisms via Population Pharmacodynamic Modeling. PLoS Comput Biol. 2016 Mar 1;12(3). doi:10.1371/JOURNAL.PCBI.1004782 PubMed PMID: 26967893.

8. Mi K, Zhou K, Sun L, Hou Y, Ma W, Xu X, et al. Application of Semi-Mechanistic Pharmacokinetic and Pharmacodynamic Model in Antimicrobial Resistance. Pharmaceutics 2022, Vol 14, Page 246. 2022 Jan 21;14(2):246. doi:10.3390/PHARMACEUTICS14020246

9. Vaisbourd E, Glass DS, Yang Y, Mayo A, Bren A, Alon U. Principles of Bacteriostatic and Bactericidal Antibiotics at Subinhibitory Concentrations. mBio. 2025 Nov 12;16(11):1–17. doi:10.1128/MBIO.02066-25;WEBSITE:WEBSITE:ASMJ;ISSUE:ISSUE:DOI PubMed PMID: 41104949.

10. Wen X, Gehring R, Stallbaumer A, Riviere JE, Volkova V V. Limitations of MIC as sole metric of pharmacodynamic response across the range of antimicrobial susceptibilities within a single bacterial species. Scientific Reports 2016 6:1. 2016 Dec 1;6(1):37907-. doi:10.1038/srep37907 PubMed PMID: 27905408.

11. Nielsen EI, Cars O, Friberg LE. Predicting in vitro antibacterial efficacy across experimental designs with a semimechanistic pharmacokinetic-pharmacodynamic model. Antimicrob Agents Chemother. 2011 Apr;55(4):1571–9. doi:10.1128/AAC.01286-10 PubMed PMID: 21282424.

12. Galvanin F, Ballan CC, Barolo M, Bezzo F. A general model-based design of experiments approach to achieve practical identifiability of pharmacokinetic and pharmacodynamic models. J Pharmacokinet Pharmacodyn. 2013 Aug;40(4):451–67. doi:10.1007/S10928-013-9321-5 PubMed PMID: 23733369.

13. Janzén DLI, Bergenholm L, Jirstrand M, Parkinson J, Yates J, Evans ND, et al. Parameter Identifiability of Fundamental Pharmacodynamic Models. Front Physiol. 2016;7(DEC). doi:10.3389/FPHYS.2016.00590 PubMed PMID: 27994553.

14. Rios ACA, Pham AD, Knibbe CAJ, Rozen DE, Hasselt JGC van, Aulin LBS. The influence of heteroresistance, growth and antibiotic selection in shaping the invasion dynamics of colistin resistant Pseudomonas aeruginosa. *bioRxiv*. 2025 Dec 15;2025.12.15.694340. doi:10.64898/2025.12.15.694340

15. Zeileis A, Leisch F, Hornik K, Kleiber C. Testing, Monitoring, and Dating Structural Changes [R package strucchange version 1.5-4]. CRAN: Contributed Packages. 2024 Sep 2. doi:10.32614/CRAN.PACKAGE.STRUCCHANGE

16. Santiago M, Hurtado I. Package “BreakPoints” Type Package Title Identify Breakpoints in Series of Data. 2026. doi:10.1002/joc.3370060607

17. Muggeo V. Segmented: An R Package to Fit Regression Models With Broken-Line Relationships. R News. 2008 Jul;8:20–5.

18 R Core Team. R: The R Project for Statistical Computing [Internet]. 2026 [cited 2026 Jul 27]. Available from: https://www.r-project.org/

19. Fidler M, Wilkins JJ, Hooijmaijers R, Post TM, Schoemaker R, Trame MN, et al. Nonlinear Mixed-Effects Model Development and Simulation Using nlmi:r and Related R Open-Source Packages. CPT Pharmacometrics Syst Pharmacol. 2019 Sep 1;8(9):621–33. doi:10.1002/PSP4.12445;REǪUESTEDJOURNAL:JOURNAL:21638306;WGROUP:STRING: PUBLICATION PubMed PMID: 31207186.

20. Sher A, Niederer SA, Mirams GR, Kirpichnikova A, Allen R, Pathmanathan P, et al. A Ǫuantitative Systems Pharmacology Perspective on the Importance of Parameter Identifiability. Bulletin of Mathematical Biology 2022 84:3. 2022 Feb 7;84(3):39-. doi:10.1007/S11538-021-00982-5 PubMed PMID: 35132487.

21. Galvanin F, Ballan CC, Barolo M, Bezzo F. A general model-based design of experiments approach to achieve practical identifiability of pharmacokinetic and pharmacodynamic models. Journal of Pharmacokinetics and Pharmacodynamics 2013 40:4. 2013 Jun 4;40(4):451–67. doi:10.1007/S10928-013-9321-5 PubMed PMID: 23733369.

22. Dunne A. An iterative curve stripping technique for pharmacokinetic parameter estimation. J Pharm Pharmacol. 1986;38(2):97–101. doi:10.1111/J.2042-7158.1986.TB04519.X PubMed PMID: 2870170.

23. Chung P, McNamara PJ, Campion JJ, Evans ME. Mechanism-based pharmacodynamic models of fluoroquinolone resistance in Staphylococcus aureus. Antimicrob Agents Chemother. 2006 Sep;50(9):2957–65. doi:10.1128/AAC.00736-05 PubMed PMID: 16940088.

24. van Os W, Pham AD, Eberl S, Minichmayr IK, van Hasselt JGC, Zeitlinger M. Integrative model-based comparison of target site-specific antimicrobial effects: A case study with ceftaroline and lefamulin. Int J Antimicrob Agents. 2024 May 1;63(5):107148. doi:10.1016/J.IJANTIMICAG.2024.107148 PubMed PMID: 38508535.

25. van Os W, Nussbaumer-Pröll A, Pham AD, Wijnant GJ, Pokem PN, Van Bambeke F, et al. Pharmacokinetic/pharmacodynamic model-based optimization of temocillin dosing strategies for the treatment of systemic infections. J Antimicrob Chemother. 2024 Oct 1;79(10):2484–92. doi:10.1093/JAC/DKAE243 PubMed PMID: 39030832.

26. Alarcon Rios AC, Zwep LB, Vriesendorp B, Knibbe CAJ, Ciofu O, Liakopoulos A, et al. Colistin resistance dynamics in Pseudomonas aeruginosa under biofilm and planktonic growth. Antimicrob Agents Chemother. 2025 Oct 1;69(10). doi:10.1128/AAC.00421-25;JOURNAL:JOURNAL:AAC;WGROUP:STRING:PUBLICATION PubMed PMID: 40900213.

