## Supplementary material for "Segmented regression to streamline semi-mechanistic pharmacodynamic modelling of antibiotic effects"

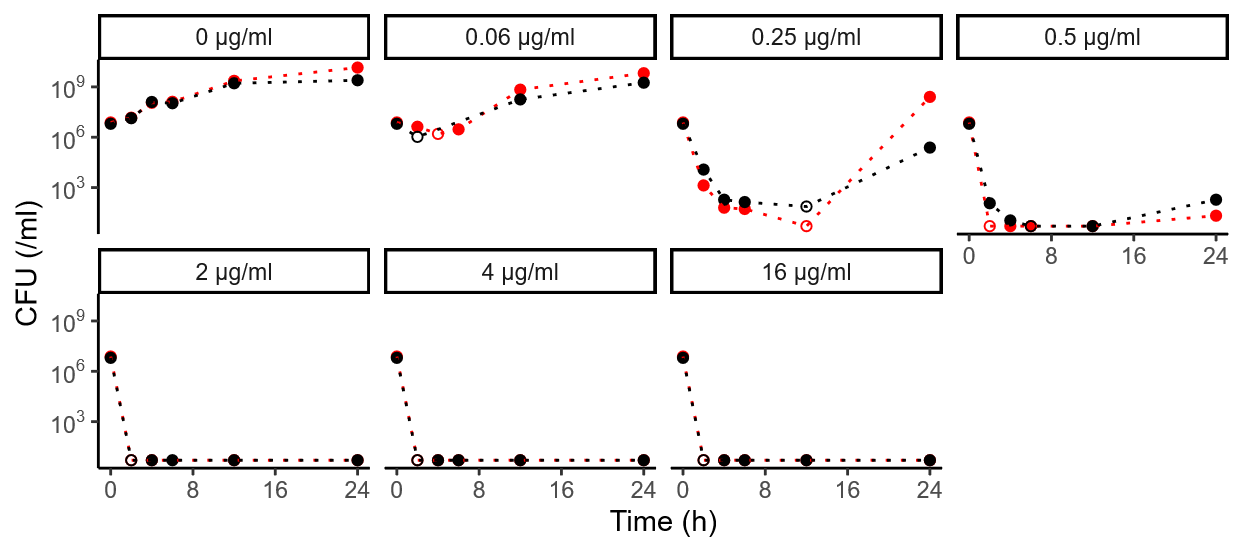

**Figure S1.** Visualisation for the time-kill data of isolate PA11976 experiments (individual facet) over time (x-axis) with CFU measurements (y-axis), with minimum CFU count used for phase segmentation presented as open circles. Replicates are indicated by colour
